# Proximity and current alignment drives fertilisation success in a broadcast-spawning coral

**DOI:** 10.1101/2025.05.24.654664

**Authors:** Gerard F. Ricardo, Christopher Doropoulos, Adriana Humanes, Liam Lachs, Helios M. Martinez, Greta Sartori, Taison Ka Tai Chang, Apple Pui Yi Chui, Elvis Long Ching Wong, Jamie R. Stevens, Iva Popovic, James R. Guest, Elizabeth Buccheri, David Idip, Peter J. Mumby

## Abstract

As coral populations decline under climate change and other stressors, surviving corals will become increasingly isolated. While Allee effects that reduce gamete encounters and fertilisation success may lead to reproductive and recruitment failure, the critical population densities and distances between individuals required to maintain viable populations remains uncertain. Here, we investigate the links between colony isolation and fertilisation patterns using an experimentally manipulated patch of the broadcast spawning tabular coral, Acropora hyacinthus. Corals were arranged in a clustered radial design, with colonies arranged at increasing distances radiating downstream from a central spawning aggregation. Fertilisation and paternity assignment analyses were used to examine the influence of parental distance, current alignment, colony size, and genetic relatedness. Fertilisation success declined sharply with increasing distance among corals, and paternity assignments indicated most parents were located within 3 m of each other, underscoring the importance of colony proximity. Additionally, fertilisation success was highest for colonies positioned downstream of the central cluster, and 84% of sequenced progeny were sired by colonies located directly upstream. Simulations of natural reef spatial population distributions projected onto a virtual grid, incorporating empirically derived parentage distances, indicated that populations remained well-mixed at typical adult densities in nearby reefs. However, as densities decreased to 1 colony per 100 m^-2^ (i.e. 0.01 colonieslllm^-2^), reproductive isolation and the formation of patchy breeding units became prevalent. Overall, our findings highlight that the spatial arrangement and isolation of corals on reefs, and subsequent Allee effects on fertilisation, can jeopardise the reproductive success of broadcast spawning corals. Conservation efforts are needed to help maintain viable coral populations and successful reproductive aggregations for degraded reef systems.

## Introduction

Allee effects are a density-dependent phenomenon where reductions in population size and density lead to diminishing individual fitness (e.g., survival, reproduction) and slower population growth rates, risking the possibility of local extinctions (Courchamp et al. 1999, Kramer et al. 2009). In coral reef ecosystems increasingly threatened by climate change, a significant proportion of reefs are now expected to become degraded to a point at which the size and density of populations could become insufficient for successful reproduction (Knowlton 2001, Frieler et al. 2013, IPCC 2018). Such critical Allee effects are especially pertinent during the external fertilisation process in sessile species that engage in broadcast spawning, where the success of fertilisation is dependent on the quantity and concentration of gametes produced (Levitan & Petersen 1995, Yund 2000). The proximity of conspecific spawning colonies and individual factors like colony size are known to influence gamete output and concentration (Levitan et al. 2004, Álvarez-Noriega et al. 2016, Mumby et al. 2024). These high concentrations are crucial for eggs and sperm to encounter each other for fertilisation during a brief window of a few hours, before wind and currents disperse the gametes (Oliver & Babcock 1992, Coffroth & Lasker 1998, Teo & Todd 2018). When population sizes decline, both the total reproductive output (i.e., the number of eggs, which sets the upper limit for the number of possible embryos) and the proportion of these that are successfully fertilised decrease. If severe, fertilisation failure can lead to recruitment failure for the entire reproductive season (Harrison et al. 1984), adversely affecting the ecosystem’s long-term ability to withstand the challenges posed by ocean warming and other stressors (Svensson et al. 2005, Mumby & Anthony 2015). Further, recent delineations of cryptic species through genomic analyses suggest that breeding coral populations are at lower densities than earlier estimates (Sheets et al. 2018, Grupstra et al. 2024, Riginos et al. 2024), and such that some coral species may be closer to critical population thresholds than previously thought.

In the dynamic environment of a coral spawning event, where eggs and sperm are suspended in the upper water column, gametic mixing is shaped by a range of abiotic and biotic factors that ultimately influence fertilisation success (Crimaldi 2012). In theory, heterogeneity in colony sizes, their spatial arrangement and the directional current of water all contribute to a complex mosaic of conditions that can either facilitate or hinder gamete encounters (Coffroth & Lasker 1998). Larger individuals, with their greater gamete output, may dominate fertilisation opportunities through increased egg-sperm encounters and gametic mixing. Moreover, the directionality and strength of water currents play a pivotal role in shaping the fate of the spawn slicks and suspensions (Coffroth & Lasker 1998, Gouezo et al. 2025). Currents typically disperse gamete concentrations, decreasing the likelihood of successful fertilisation. However, some physical and hydrodynamic features, such as fronts, eddies, and convergence zones, can contribute to the formation of dense ‘slicks’ of embryos in the hours to days following spawning (Oliver & Willis 1987, Wolanski & Hamner 1988, Doropoulos et al. 2019, Wolanski et al. 2024). While these features can concentrate embryos and decomposing reproductive material, their role in enhancing fertilisation remains unclear, as these hydrodynamic-driven aggregations may occur too late to significantly influence gamete encounters.

A key opportunity in advancing our understanding of fertilisation patterns lies in the incorporation of parent–offspring relationship data alongside the strategic isolation of targeted colonies from conspecifics. While most studies of fertilisation success in corals and other marine invertebrates rely on statistical associations with biophysical and spatial population parameters, they often fall short of establishing direct causal links. For instance, although variations in spatial population parameters (e.g. colony density) may correlate with fertilisation outcomes, without parental lineage data and controlled spatial context, it remains unclear whether these factors actively drive the observed patterns. Additionally, samples taken from spawn slicks to estimate fertilisation success – especially those collected the day after spawning – are prone to biases. This is because unfertilised eggs decompose, resulting in underestimates of initial egg counts and, consequently inflated fertilisation rate estimates (Oliver & Willis 1987). Even when sampling occurs within the fertilisation time window, it can remain challenging to determine whether embryos and their fertilising sperm originated from colonies within the area of interest or nearby reefs (Oliver & Willis 1987). There is also the possibility that embryos are the result of hybridisation between closely related species, complicating intraspecific fertilisation interpretations and raising uncertainty about whether these hybrid embryos will develop successfully beyond the embryo stage (Willis et al. 1997, Vollmer & Palumbi 2002). Parental assignment methods (e.g. progeny arrays) allow direct characterisation of fertilisation patterns by linking offspring to specific adult colonies. Coupled with spatial data, they offer direct insights into how population characteristics determine reproductive success (Warner et al. 2016, Dubé et al. 2020, Ricardo et al. 2024a).

Typically, at any given reef, broadcast spawning occurs over one or two nights annually, within a narrow timeframe of 2–3 hours before the cloud of gametes disperses beyond the threshold for effective fertilisation (Oliver & Babcock 1992, Omori et al. 2001, Kitanobo et al. 2022). Although laboratory-based research has shed much light on species-specific fertilisation mechanisms (Willis et al. 1997, Nozawa et al. 2015, dela Cruz & Harrison 2020, Buccheri et al. 2023), such studies generally ignore the crucial hydrodynamic influences acting on a natural spawn concentration. As such, we know little about how species-specific spawn dynamics relate to spatial population parameters in situ (but see Coma and Lasker (1997), Levitan et al. (2004), Miller and Mundy (2005), Mumby et al. (2024)). Understanding thresholds in these spatial parameters is not only critical for predicting the reproductive viability of natural reef populations under degradation, but also has direct implications for applied conservation. In particular, restorative approaches such as coral outplanting rely on assumptions about effective fertilisation, yet key knowledge gaps remain regarding how colony spacing and spatial arrangement influence in situ fertilisation rates and, consequently, the long-term contribution of outplanted colonies (Ladd et al. 2019, Suggett et al. 2019, Boström-Einarsson et al. 2020). Gathering empirical data during these annual spawning events presents significant challenges due to their ephemeral nature, nocturnal timing, numerous unknown phenological cues, and the complexity introduced by the simultaneous spawning of multiple species. Here, we aim to address these shortcomings by conducting a manipulative field experiment in which we positioned adult colonies in a radial arrangement at varying distances from a high-density central spawning hub to assess how colony spacing and arrangement influence fertilisation success and potential Allee effects. We utilise paternity assignments based on next-generation sequencing of adult colonies and individual larval progeny from the generated spawn plume to understand how gametic mixing, colony distance, directionality, colony size and genetic relatedness influence fertilisation outcomes.

## Methods

### Coral collection and experimental site selection

Here we assessed fertilisation success of the reef-building table coral Acropora hyacinthus in Palau, Micronesia. Acropora hyacinthus is a large hermaphroditic simultaneous broadcast spawning coral found in a range of physical environments and its fast growth allows it to facilitate reef recovery (Ortiz et al. 2021). Most corals including A. hyacinthus spawned on the inshore reefs adjacent to our reef site (see below) on the 1 April 2023, five days before the full moon, coinciding with warm doldrums (∼33 °C water temperature). On the 2 April 2023, approximately 95% of colonies of A. hyacinthus (Dana, 1846) were gravid at the offshore Uchelbeluu Reef (7.2591° N, 134.5295° E), and 25 medium-sized colonies of unique genotypes were subsequently collected.

To ensure a controlled experiment, a field site was selected in a sandy seagrass area at the entrance to Ngermid Bay (also known as Nikko Bay, 7.3135° N, 134.4960° E) (Fig. 1a), a complex of channels and bays on the southeastern side of Koror, Palau, known for its diverse coral communities (Golbuu et al. 2016). This site was chosen because it was free of natural populations of our target species, Acropora hyacinthus (see below). While Ngermid Bay does host a variety of corals, including Porites (56%) and Goniopora (9%), but relatively few Acropora are present (∼1%) (Kurihara et al. 2021). Notably, because the inshore Acropora colonies spawned earlier (April 1st) than our experimental patch of A. hyacinthus, and this species was not observed within the bay, we concluded that the risk of gamete contamination from these inshore colonies was unlikely. Although natural populations of the genus Acropora exist outside Ngermid Bay (Kurihara et al. 2021), we assumed that any Acropora sperm released after April 1st would be carried offshore by tidal currents, effectively maintaining spatial separation from our experimental gametes and minimising the likelihood of hybridisation (Furukawa et al. 2024).

**Fig. 1.**
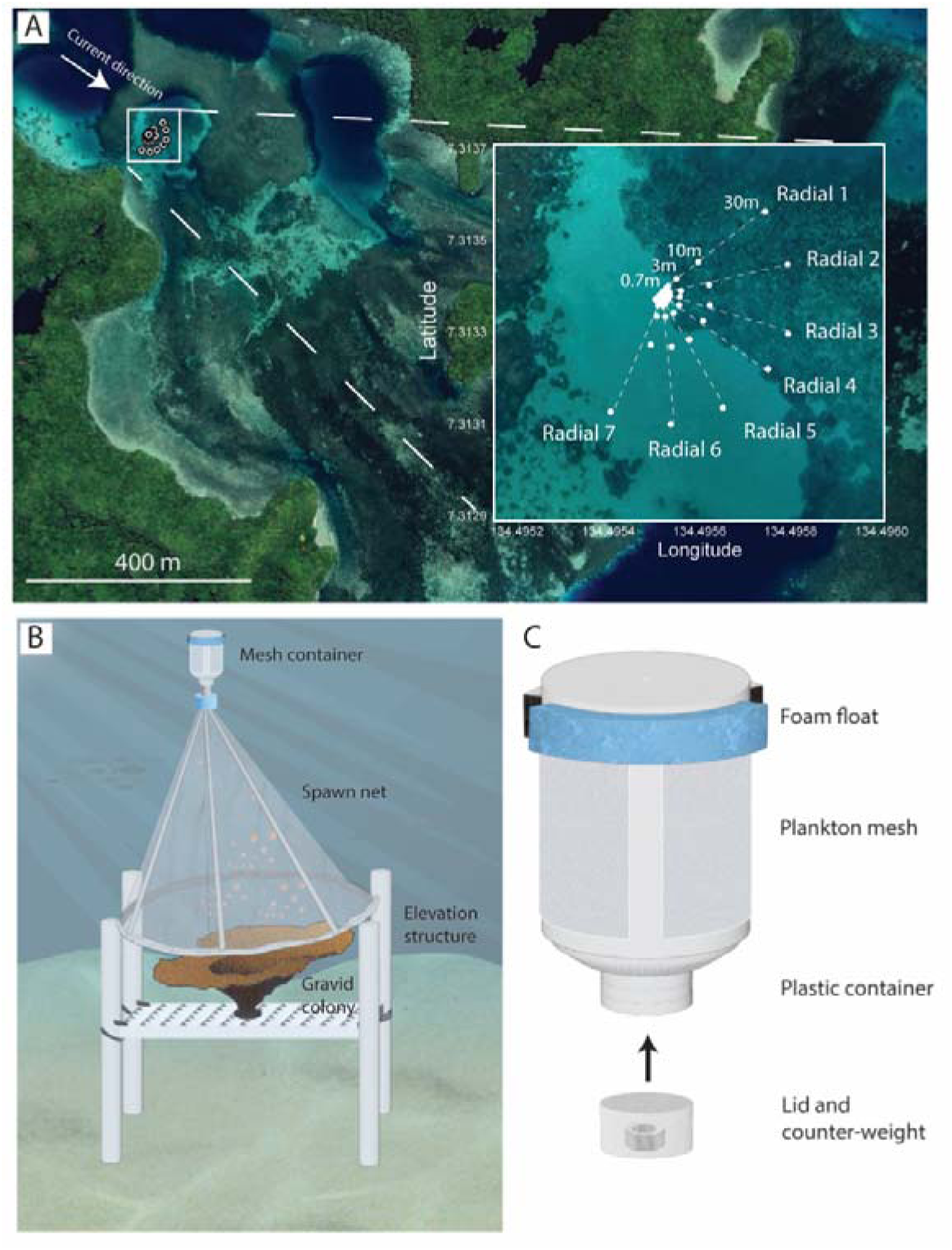
Experimental design and coral spawning catcher apparatus. (A) The clustered radial arrangement of adult gravid colonies of Acropora hyacinthus, with colonies positioned on the radial lines at distances of 0.7, 3, 10, 30 m from the central patch. (B) Design diagrams of spawning structures used to catch and funnel the egg-sperm bundles, and (C) the design of the mesh containers that retained the adult colony’s eggs and were released into the concentration of gametes produced from the experimental patch.

### Experimental design

The experiment was designed to assess the effect of intercolonial spacing and spatial arrangement of spawning colonies on fertilisation success, utilising several radial lines extending from a central high-density patch of gravid corals in a semi-circle shape (Fig. 1a). The experimental arrangement was strategically oriented such that the central radial extensions were positioned downstream relative to a high-density aggregation of colonies, aligning with the anticipated tidal cycles and current flows characteristic of the bay. Seventeen A. hyacinthus colonies (of 16 unique genotypes) were arranged in a semi-circle, each spaced 0.7 m apart measured from the centre of each colony. Extending from this semi-circle, seven radial lines were marked, with reference points for coral placement at 0.7, 3, 10, and 30 m intervals along axes set at 0°, 26°, 52°, 78°, 104°, 130°, and 156° relative to the perpendicular of the anticipated current direction (Fig. 1a). To avoid potential smothering of sand, coral colonies were elevated approximately 30 cm above the seabed using structures composed of 0.7 m pipes and egg crate material, all of which were georeferenced using handheld GNSS receivers (Fig. 1a). To mitigate the risk of cross-fertilisation between adjacent colonies within a radial line, larger colonies were fragmented and strategically distributed across designated positions along that colony’s corresponding radial line. This arrangement aimed to minimising the potential for cross-fertilisation among the individual colonies positioned along these radial lines so maximum colony distances could be assessed with little interference. The average diameter of the coral colonies used in the experiment was 32.6 ± 12.9 cm, slightly larger than the 26.0 ± 13.1 cm diameter observed in natural populations on a nearby offshore reef (Ricardo et al. 2024b).

Immediately prior to coral spawning, coral spawn catchers were positioned approximately 20 cm above the colonies along the radial lines 2 to 7 (Fig. 1b), as well as above four randomly selected central colonies. The spawn catchers channelled the egg-sperm bundles into detachable 270 mL containers. Approximately 46% of each container’s wall was removed and replaced with a 250 µm mesh, allowing sperm to diffuse freely while retaining the eggs of individual colonies (herein referred to as ‘mesh containers’). As a result, eggs from tracked colonies (hereafter referred to as egg donors) were fertilised by sperm from initially unidentified colonies (hereafter referred to as fertilising sperm donors). While all in situ fertilisation collection methods introduce some degree of artefact (see review by Levitan (1995)), egg containment is often necessary in studies of small populations where natural egg retrieval via plankton sampling is infeasible or where mixing with eggs from other species is a concern. To facilitate tracking, glowsticks were affixed to the mesh containers using cable ties. Each container was modified with flotation on the top and a bolt counterweight in the base, ensuring it remained vertically oriented with ∼10 cm submerged in the water column (Fig. 1c).

During the predicted coral spawning window, divers entered the water at ∼18:00 and monitored corals along the patch using red beam dive torches. All but one monitored colony commenced spawning within a 12 min period of each other. Once individual colonies were spawning, the mesh containers were removed from the catcher, capped, the glowsticks activated, and containers released. Due to logistical constraints, retrieval was conducted via kayak in shallow areas inaccessible by boat approximately 80–90 min after release. Following collection, samples remained submerged in the ambient water post-collection until they were processed in the laboratory. While this prolonged gamete contact time, any additional fertilisation resulting from extended exposure is assumed to be negligible given rapid sperm dilution rates (Omori et al. 2001).

On the night of the experiment, the tide was at mid-ebb (0.67 amplitude), flowing SSE toward the bay entrance. The site was largely sheltered from the prevailing 2.4-knot NNW offshore wind, resulting in the gamete concentrations and mesh containers being predominantly driven by tidal currents. Water depth downstream of the experimental setup varied between 1 and 3 m, with mesh containers drifting at a mean velocity of 0.21 m s^-1^ (n = 3), placing them in the 97th percentile of water speeds recorded by the Marotte tilt-meter during a two-day deployment under a comparable tidal regime. Additionally, tracer dye (fluorescein) releases were conducted at the site following Ricardo et al. (2024b) during comparable tidal cycles as the main experiment using an Unmanned Autonomous Vehicle (UAV: DJI Phantom IV) to understand the movement and dispersion properties likely acting on the gamete plume.

### Fertilisation success

Samples were fixed with 4% buffered formalin in filtered seawater containing 10 g L^−1^ sodium β-glycerophosphate at a ratio of 1:4 fixative to sample, approximately ∼150 min after spawning when the developing embryos reached the four-cell cleavage stage. Samples were later assessed under a dissection microscope for fertilisation success, recorded as the number of cleaved embryos as a proportion of total initial eggs.

A number of controls were conducted on a nearby vessel shortly after the release of the mesh containers, by attaching closed containers to spawn catchers of randomly selected colonies before transporting them to the vessel. First, two separate one-to-one crosses were conducted from within the experiment which resulted in 71–80% fertilisation success. Next, two self-fertilisation controls were conducted by capturing released bundles from two colonies within the experimental patch in closed containers. One container resulted in unrealistically high self-fertilisation (18.1%) in contrast to the other sample (1.0%), the paternity assignments (below), and previous self-fertilisation reported of this species in Palau (0.1%) (Ricardo et al. 2024b), suggesting that this container was likely contaminated during the collection processes, as it was uncapped and attached to the spawn catcher. Finally, although not tested in this study, previous work has shown that sperm can move freely through the plankton mesh of the containers with resulting fertilisation success within the range of their mesh-free control (Mumby et al. 2024, Ricardo et al. 2024b).

### Genetic sampling and sequencing

A small amount of tissue from each of the adult colonies was collected into individual ziplock bags. The tissue for each individual was further divided into two and placed in 1 mL of 100% ethanol in a 2 mL Eppendorf and placed in a -80 °C freezer. To assess paternity, genetic tests were conducted on individual larvae allowing determination of the fertilising sperm donors from the sample from known egg donor colonies following Ricardo et al. (2024a). Fertilised eggs from the mesh containers were separated individually and allowed to develop until 36–42 h old and then pipetted with 10 µL seawater in 100 µL of 100% Ethanol. After 24 hr, a 50% ethanol exchange was conducted on each sample to further remove seawater and contaminants. Samples were shipped back to the University of Queensland, Australia (CITES permit number PW23-079).

A total of 82 adult colony (n = 2 per colony) and 139 larval samples were transferred to 96-well plates (Eppendorf, TwinTec©, skirted, 150 μL, PCR clean), capped (Eppendorf, Cap Strips, 8-strips, domed), and sent to Diversity Arrays Technology (DArTSeq; Canberra, Australia) for DNA extraction, library preparation, sequencing and single nucleotide polymorphisms (SNP) identification. DArTSeq sequencing is a technique that employs pairs of restriction enzymes to digest genomic DNA, similar to other site-associated reduced representation methods (e.g., RADseq) enabling the identification of genome-wide SNP markers. Sequencing was carried out on an Illumina short-read platform to produce raw “sequence tags” of approximately 75 base pairs. Samples were aligned to two reference genomes for Acropora hyacinthus: Acropora_hyacinthus_chrsv1 (accession number: GCA_020536085.1) and Acropora_hyacinthus_v002 (available at http://ahya.reefgenomics.org/). The raw sequence reads were processed following DArT’s proprietary variant calling pipeline. Raw reads were filtered based on sequence quality, especially in the barcode region, truncated to 69 base pairs, and grouped by sequence similarity. Subsequently, a series of filters was applied to select sequence tags that contain reliable SNP markers. One-third of the samples were processed twice as technical replicates, starting from DNA and using independent adaptors, through to allelic calls. Then, a series of proprietary filters were applied to select the sequence tags that contain reliable variants. High-quality markers with low error rates were primarily selected using scoring consistency (repeatability), assessed using technical replicates.

Raw SNP sequencing data underwent standard filtering processes for paternity assignments following (Premachandra et al. 2019, Jenkins et al. 2024). First, monomorphic loci were removed, resulting in 50,405 SNPs. Next, an initial filtering step was applied to remove loci > 50% missing data and to exclude loci with a minor allele frequency (MAF) <0.05 to minimise the possibility of scoring sequencing errors as rare minor alleles. Loci were then further filtered based on read depth at a minimum mean threshold of >3 reads. A stricter filtering step was then applied to retain only loci with a call rate >95% and a mean read depth of >20, followed by removal of loci with excessively high read depth based on a three-standard-deviation range. Secondary loci were also eliminated using a random selection method, and loci were assessed for reproducibility, with those <95% threshold discarded. Finally, loci deviating significantly from Hardy-Weinberg equilibrium (HWE) were filtered and false discovery rate (FDR) multiple testing corrections applied. The final dataset consisted of 1,328 SNPs across 81 adult samples and 120 larval samples, which was used in downstream analyses. Prior to paternity assignments, final filtering was conducted following Premachandra et al. (2019) to remove mean MAF values <0.2. Typically, only low numbers of high-quality SNPs are required for accurate parentage assignments (Anderson & Garza 2006, Premachandra et al. 2019).

#### Paternity assignments

Paternity assignments were conducted on A. hyacinthus larval samples with known maternal genotypes at the time of egg capture. Paternity assignments were carried out using the software CERVUS 3.0.7 (Kalinowski et al. 2007). CERVUS is a simulation-based approach that uses likelihood ratios to assign parentage. The simulation-based approach is relatively robust to null alleles and genotyping errors. Simulations were based on allele frequencies of the experiment population; 20,000 offspring, a 5% error rate, and 95% sampled candidate fathers were used as inputs. A 95% ‘sampled candidate fathers’ rate was chosen to account for a few individuals that were not successfully sequenced. Since CERVUS cannot identify unsampled candidate parents during assignments, this uncertainty is reflected in the confidence levels of the assignments. The assignments were conducted under the ‘strict’ (95%) confidence levels using log-likelihood ratio (LOD) scores. Assignments for larvae were removed when the sperm donor could result from multiple positions i.e. from fragments along the radial lines.

#### Population structure

Clones within the sample set, including fragments used within the radial lines, were assessed using COLONY v2.0.7.1 (Jones & Wang 2010), a maximum likelihood-based software designed for inferring sibship, parentage, and clonemates from multilocus genotypic data. The analysis was run on 81 multilocus genotypes using the full-likelihood (FL) method, with medium run length and high precision settings. The dataset was specified as monoecious, diploid, with clonal inference enabled. Clone assignment in COLONY is achieved via a two-step process: first, full-sib families are inferred, and then clone groups are delineated within these families by maximising the likelihood of shared multilocus genotypes, accounting for potential mistyping. All clone groups were identified with high confidence (most with posterior probabilities of 1.000). Clones and replicates were removed before conducting the population structure analysis, ensuring that only unique genotypes were compared.

To assess the population genetic structure within the population, we performed Principal Component Analysis (PCA) on quality-filtered allele frequency data using the package ade4 (Dray & Dufour 2007), in R (v 4.4.1). PCA was conducted with centring and scaling, and the first 18 principal components were analysed which represented ∼90% of the variance. To test for population clustering, we applied k-means clustering, and the number of clusters was assessed using the Elbow Method (total within-cluster sum of squares) and Silhouette scores. Additionally, we performed STRUCTURE analysis within the package dartR, using an admixture model with correlated allele frequencies. STRUCTURE was run with a burn-in of 10,000 iterations, followed by 50,000 MCMC replications, testing K = 1 to K = 5, with two replicates per K. The optimal K was determined using the Evanno method (Evanno et al. 2005). To further assess genetic differentiation, we conducted a PERMANOVA in the package vegan (Oksanen et al. 2007), testing whether genetic distances differed significantly between groups, with 9,999 permutations to evaluate statistical significance.

### Statistical analysis

Where necessary, fertilisation success data were subsampled to balance differences in sample sizes between containers. The relationship between fertilisation success with degree deviation (i.e., 0°) from directly downstream of the central patch (angle, continuous, modelled non-linearly) and distance from the central patch (dist_center, continuous) was analysed using a Generalized Additive Mixed Model (GAMM) with a binomial family:

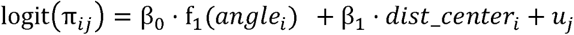

where u_j_ represents an observation-level random intercept to account for overdispersion (Harrison 2015). The GAMM was implemented in R (v. 4.4.1) using the package mgcv (Wood & Wood 2015). To identify the optimal number of folds for cross-validation, k = 2 to 5 were tested and model performance (mean log-likelihood) was stable for k = 2–3, but decreased with higher k; therefore k=3 was selected for the final model. Additionally, three colonies located along the radial lines that shared a clonemate with an individual in the high-density patch were adjusted in the model to account for their relatively reduced exposure to unique sperm sources. However, including this adjustment had negligible effect on model performance and reduced parsimony, so it was not retained in the final model.

For paternity assignments, observed pairwise crosses were first analysed separately for each factor. Distances between assigned parents were calculated with and without weighting using:

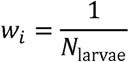

where *N*_larvae_ was the number of larvae per container. Weighting was applied to prevent an overrepresentation of highly sequenced containers, though this approach can disproportionately amplify trends in containers with low sample sizes. A Kolmogorov-Smirnov (K-S) test was used to assess if the pairwise-cross distance distribution differed from random chance based on the distance distribution between candidate adults. Additionally, a Rayleigh’s test for uniformity was conducted to assess whether the spatial distribution of colonies relative to the current direction deviated from uniformity.

Next, the statistical relationships between individual or population parameters and assigned parent colonies were examined. Observed pairwise combinations of parents were treated as count data and compared against every possible pairwise combination of candidate adults in the experiment, excluding unlikely combinations such as those between known genotype fragments. Given the combination of true and false zeros – where total embryos in a container after fertilisation exceeded those larvae eventually sequenced – in the dataset (Martin et al. 2005), relationships with predictor variables were assessed using zero-inflated negative binomial model using the package glmmTMB (Magnusson et al. 2017). To account for excess zeros, including both true and false zeros, the zero-inflation component was tested using three approaches: (1) total embryos per container as a predictor, (2) a constant (intercept-only model), and (3) without the zero-inflation component altogether. Treatments included distance between parent colonies (distance), angle of parent colonies relative to upstream/downstream direction (angle), colony size (col_size) and genetic relatedness (gen_dist). We hypothesised: (1) a negative relationship between colony distance and fertilisation success, moderated by colony angle, (2) a positive effect of colony size, and (3) a non-linear (quadratic) effect of genetic distance, peaking at intermediate values. To address collinearity, covariates were centred and scaled, resulting in a Variance Inflation Factor (VIF) <2 for all predictors.

A global model was constructed with biologically meaningful interactions and factors, incorporating an offset term for total fertilisation in each mesh container. The offset helped mitigate biases introduced by unequal sequencing effort across mesh containers. The model was specified as follows:

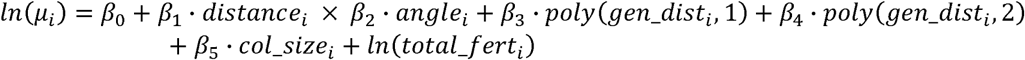

and the zero-inflation component as:

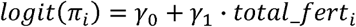

To identify the most appropriate error distribution and assess the need for a zero-inflation component, the global model was first fitted and compared across Poisson and negative binomial distributions, with and without zero-inflation, using AIC. Based on these comparisons, the most parsimonious model was a negative binomial distribution without a zero-inflation term. Model selection was then carried out using the dredge function from the MuMIn package (Barton & Barton 2015). A set of candidate models with ΔAIC < 2 were retained for model averaging. The final averaged model, based on this subset, included all original coefficients and interactions but excluded the zero-inflation term. Model assumptions were evaluated using the DHARMa package (Hartig & Hartig 2017), and there was no clear indication of model assumption violations.

### Mixing simulations

To explore the potential for gametic mixing during fertilisation within realistic population spatial structures, we simulated a 100 × 100 m virtual grid at 0.1 m resolution, populated 200 times for each of three spatial distribution types using the spatstat package. First, a random (Poisson) process was implemented. Next, we used two spatial distributions derived from natural reef populations near the study site. Belt transect data from reef crest and reef slope habitats from Ricardo et al. (2024b) were fitted to Poisson, Thomas, Matérn, and Cauchy cluster models. Based on Akaike Information Criterion (AIC), Matérn cluster models provided the best fit for both reef types. The best-fit model parameters were extracted and used to simulate colony distributions for ‘reef crest’ and ‘reef slope’ simulations. We then computed Euclidean distances for all pairwise combinations of adults within the grid. The probability of fertilisation occurring between adult pairs at least once was determined using the empirical unweighted density function from our field experiments. Simulations were run across a gradient of adult densities, ranging from low-density (highly degraded reefs, 0.0003 colonies m^-2^; Zayasu and Suzuki (2019)) to densities observed on healthy reefs (0.3 colonies m^-2^; Ricardo et al. (2024b)). The ‘number of pairwise crosses’ and ‘partners per individual’ and ‘isolated individuals’ were extracted from each run. The simulations assume that the probability of cross-fertilisation between two adult colonies is equally likely based on their distance, recognising this as a generalisation that may not fully capture all the factors influencing crossing success in nature (e.g., unidirectional current).

## Results

### Fertilisation success

Mean fertilisation success across all containers (n = 25) was 32.8%, ranging from 0 to 95%. Fertilisation success (mean ± SD) within the central patch was 50.6 ± 34.0%, whereas fertilisation in the radial lines averaged 29.0% and ranged from 0 to 95%. The samples with the highest fertilisation success, exceeding 90%, were all located directly downstream of the central patch.

Model results from the GAMM analysis revealed there were significant effects of distance and the angle of the colony from downstream current direction on fertilisation success. Specifically, distance from the centre of the patch resulted in a negative association with fertilisation success (β = -0.1168, p = 0.002), corresponding to an ∼11% decrease in the odds of fertilisation for every additional metre away from the centre, while holding ‘angle’ constant. Furthermore, the angle of the colony from downstream current direction was significant (F = 15.33, p < 0.001), indicating that fertilisation success peaked between the radial lines 5 and 6 following the main current direction (Fig. 2). Model residuals showed high variability between samples (adjusted R^2^ = 0.60), including those in close proximity, suggesting spatial patchiness. Some of this variability could be attributed to cross-fertilisation between radial lines (see paternity assignments below).

**Fig. 2.**
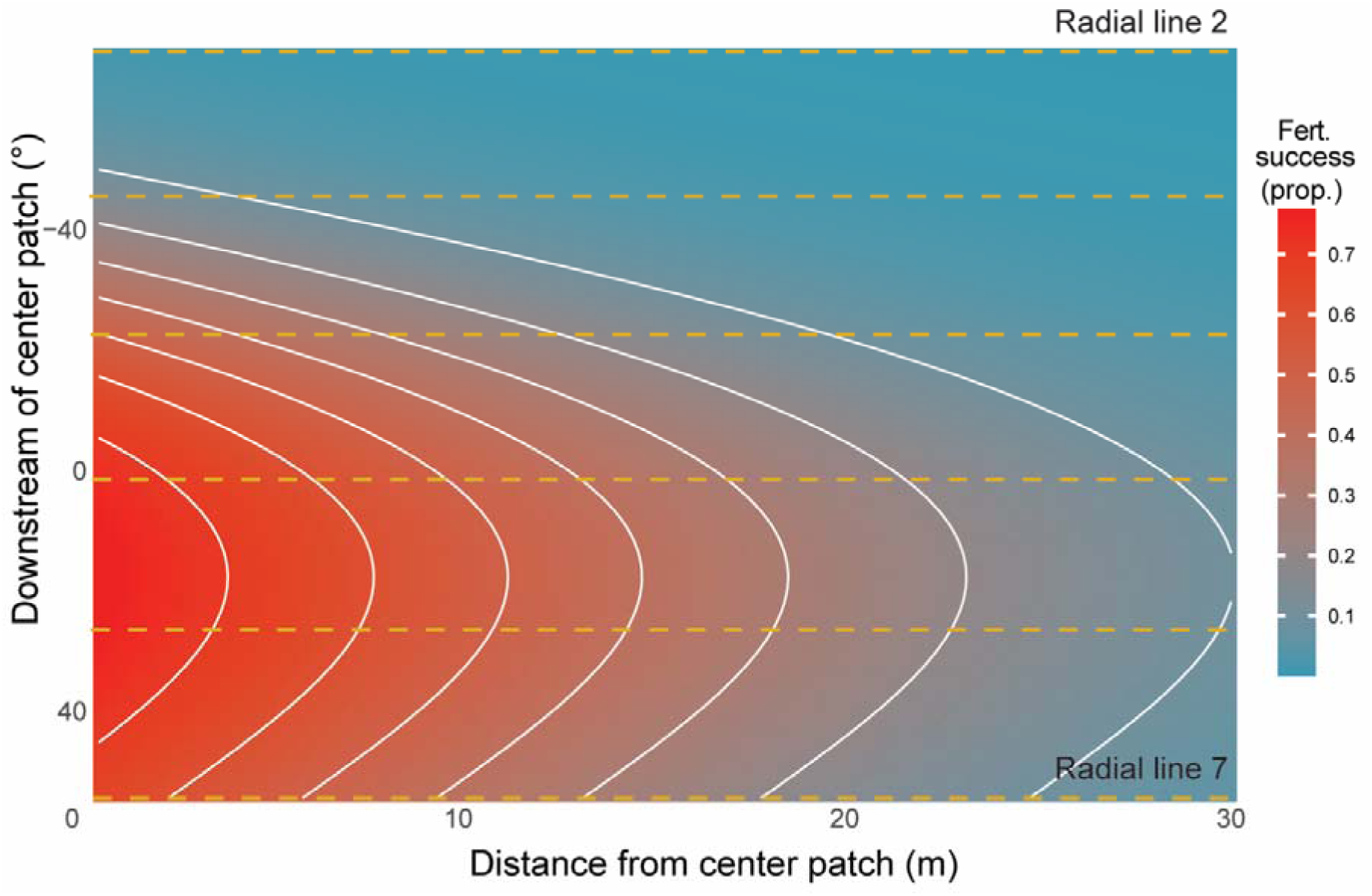
The effect of distance of colonies along each radial line relative to the central spawning patch, and their spatial position relative to the current advection line (0°) on fertilisation success (proportion fertilised), derived from the bulk fertilisation success analysis. White lines represent every 0.1 proportion interval. Orange lines denote the radiating lines in the experimental patch. Only radial lines 2 to 7 were assessed for fertilisation success.

### Paternity assignments

After excluding larvae with ambiguous paternity, a total of 62 sequenced larvae were assigned to a sperm donor with high confidence (>95%). The median distance between parental crosses was 1.8 m, whereas the median weighted distance was 2.4 m (Fig. 3a, c). When the pairwise cross distributions were compared against the distributions of the distances between individual adults, there was a significant difference (unweighted: D = 0.459, p < 0.001; weighted: D = 0.964, p < 0.001), with a preference toward lower crossing distances than expected by chance. Bearings of fertilising sperm donors relative to their respective egg donor were predominantly from 300 to 20 degrees (NW-N), aligning closely with upstream current direction (326 degrees) (Fig. 3c). A Rayleigh test of uniformity, which does not account for other covariates, indicated strong directionality of the pairwise crosses (R = 0.866, p < 0.001), with 84% of sequenced progeny sired by colonies located within 90° of the upstream direction. This directionality was supported by tracer dye releases which revealed the gametes may form longer plumes in the direction of the current under higher current velocities (Fig. S2).

**Fig. 3.**
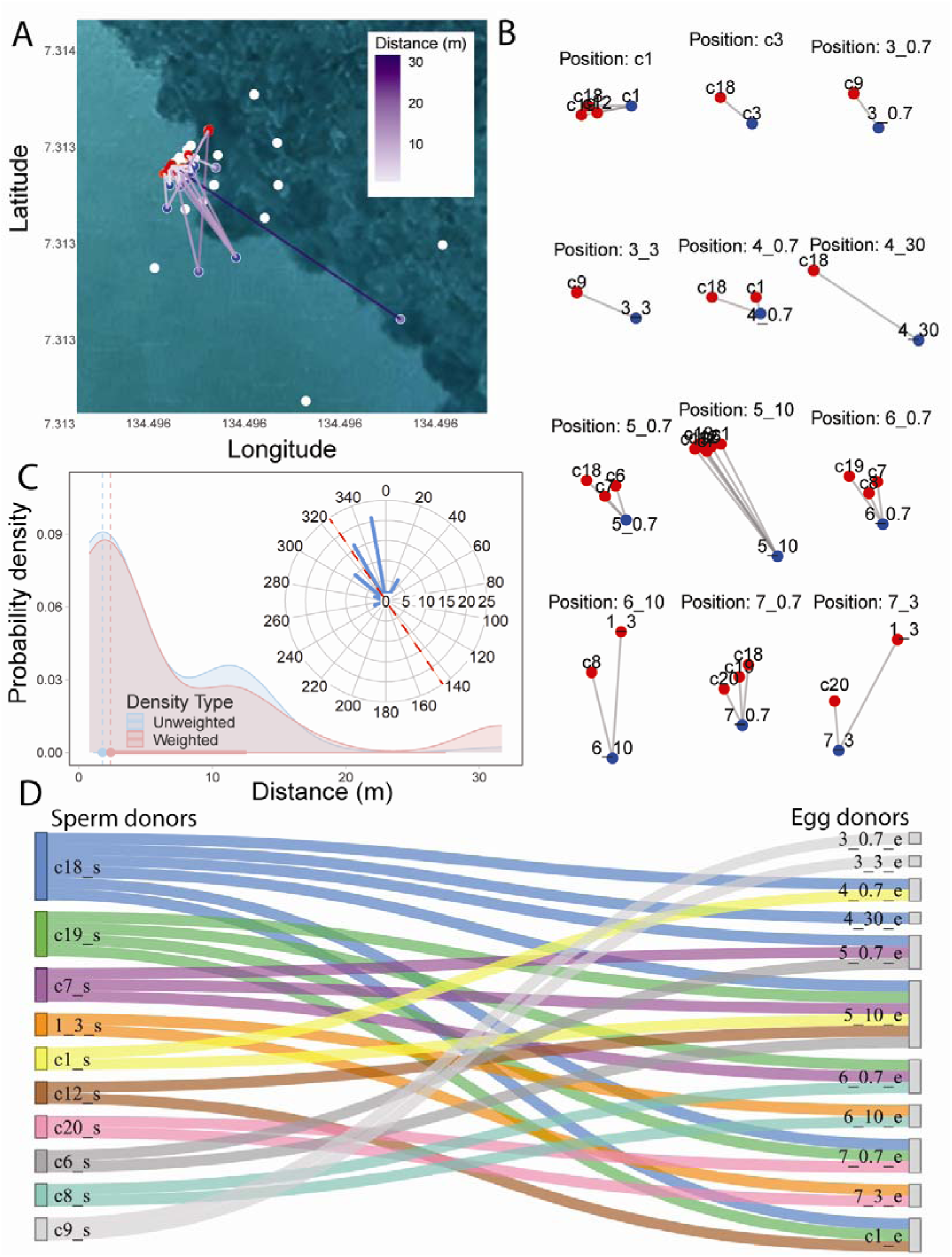
Pairwise analysis from paternity assignments. (A) All pairwise crosses within the experimental patch. Egg donors = blue nodes. Fertilising sperm donors = red nodes. A small amount of jitter was added to the sperm donors for visibility. (B) Spatial arrangement of all breeding units within the patch relative to the egg donor positions (blue). Red denotes sperm donor positions. Peripheral colonies are labelled in the format ‘RadialLine_Distance_Donor’ where is the distance from central patch, and for central patch colonies ‘cID_Donor’, where “c” indicates a central patch colony. (C) Weighted and unweighted intercolonial distances between pairwise crosses. Inset: Direction of fertilising sperm donors relative to egg donors. True north = 0. Current direction signified by the red line. (D) Sankey diagram illustrating unique pairwise crosses between identified sperm donors and known egg donor colonies. Colony labels correspond to each individual’s spatial position within the experimental design, as described above.

Not all colonies participated in cross-fertilisation; even gametes from some colonies in very close proximity sometimes failed to cross-fertilise. Among the participants, 10 colonies contributed sperm that successfully fertilised eggs from 12 colonies (Fig. 3b, d). There were 28 unique pairwise crosses, with each egg-donor colony fertilised by an average of 2.8 sperm-donor colonies. However, reproductive success was unevenly distributed such that the top three sperm donors fertilised 67% of participating egg providers at least once (Fig. 3d).

The most parsimonious statistical model describing the number of pairwise crosses included all predictors (effect plots shown in Fig. S3); with the only statistically significant predictors being distance and colony positions relative to current. The number of pairwise crosses decreased as the distance between colonies increased (β = -0.130, p = 0.004), while colonies positioned more favourably relative to current resulted in a greater number of pairwise crosses (β = 3.141, p = 0.028). Based on parental genetic relatedness, self-fertilisation was identified in five larvae, each from a different egg donor, representing 4.6% of the population. Paternity assignment also revealed noise in the bulk fertilisation analysis were largely driven by cross-fertilisation between radial lines. There was limited evidence of population structure among the collected colonies within the small spatial scale of the collection site, validating that sampled colonies belong to a single taxon (Text and Fig. S1).

### Mixing simulations

All three spatial dispersion patterns (random, reef crest and reef slope) exhibited similar trends with mixing metrics and were subsequently pooled (Fig. 4, S4). Both the mean number of pairwise crosses and the number of partners per individual followed a non-linear exponential increase with adult density (Fig. 4a, b). The inflection point for pairwise crosses occurred at 0.086 colonies m^-2^, while for colonies crossed per individual, it was 0.124 colonies m^-2^. Surprisingly, few colonies were completely isolated from mixing until densities were very low (Fig. 4c). With decreasing density, the number of isolated colonies increased – peaking at ∼0.01 colonies m^-2^ – with subsequent declines at extreme low densities where the overall number of colonies (isolated or not) becomes a limiting factor.

**Fig. 4.**
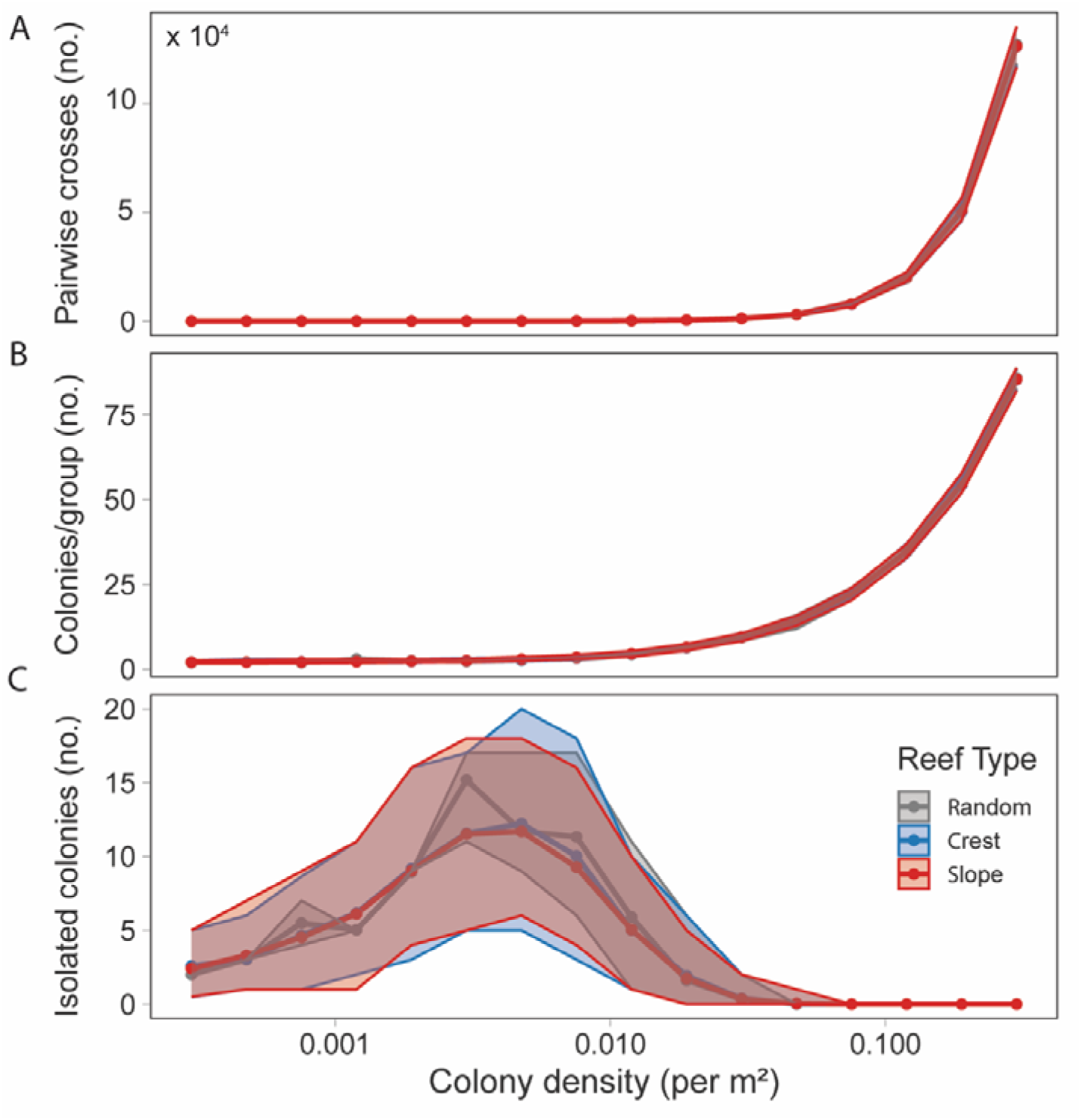
Simulated relationships between colony density and key genetic mixing metrics across different reef types in an area of 1 hectare (10,000 m²). (A) Number of pairwise crosses, (B) number of reproductive partners per individual, and (C) number of isolated individuals as a function of colony density (per m²). Shaded areas represent the 95% percentile range. Data are shown for three spatial dispersion patterns: random (grey), reef crest (blue), and reef slope (red).

## Discussion

Using a novel combination of two complementary fertilisation assessment methods with spatial manipulations of adult colonies designed to evaluate reproductive success, we reveal strong spatial constraints on coral fertilisation. We find that fertilisation in broadcast spawning corals is related to the spatial positioning of the adult colonies, particularly their alignment relative to prevailing tidal currents and their proximity to high-density spawning patches, with individuals on the periphery of the downstream current contributing less to successful fertilisation.

Paternity analyses provide critical insights into gamete interactions within the gamete cloud, capturing fertilisation information not readily apparent in bulk fertilisation estimates (Flanagan & Jones 2019). The median fertilisation distance between assigned parent pairs yielded a median fertilisation distance of 3–4 m. While this threshold is influenced by our experimental design (which included a higher density of closely spaced colonies), it aligns with predictions from biophysical models and previous empirical studies, and indicates that broadcast spawning corals isolated by beyond 20 m are limited in their potential for fertilisation (Levitan et al. 2004, Teo & Todd 2018, Mumby et al. 2024, Ricardo et al. 2024b). Surprisingly, this maximum distance of fertilisation limitation from spawning corals is similar to brooding corals that utilise internal fertilisation, despite spawner gametes often dispersing hundreds of metres during the fertilisation window. Using paternity assignments, Warner et al. (2016) found a maximum distance of 17 m for the brooding coral Seriatopora, with most fertilisation occurring within 10 m. Likewise in brooding octocorals, paternity analyses found the maximum distance of 12 m despite the relationship with proximity being less clear (Lasker et al. 2008). Our findings underscore the risk that depleted populations, with increased distances between colonies, are highly susceptible to component Allee effects that significantly impair fertilisation during spawning events.

The spatial orientation of spawning coral colonies relative to tidal currents and the presence of upstream conspecifics significantly influences fertilisation success. Under the observed conditions, where advection predominated over dispersion (hydrodynamic conditions illustrated in Fig. S2), colonies positioned downstream of one another within a patch experienced greater fertilisation success. Paternity analyses revealed that egg donor colonies were predominantly fertilised by upstream sperm donor colonies, a result consistent with previous findings (Coffroth & Lasker 1998), and attributed to current-mediated sperm transport dynamics. The high-density upstream patch of spawning corals served as the primary source of sperm for downstream egg donor colonies, supporting the unidirectional nature of fertilisation. On natural reef slopes, this alignment may arise due to topographical constraints that impose linear colony arrangements along shore-parallel tidal currents (Wolanski & Hamner 1988). However, in environments where currents shift onshore into lagoons or offshore into deeper waters under changing wind conditions, the directional advantage can be diminished and alter fertilisation success patterns (Mumby et al. 2024). Thus, the spatial arrangement of colonies relative to prevailing currents can create major fertilisation asymmetries, underscoring the importance of considering hydrodynamics in reproductive connectivity on coral reefs.

Interestingly, although the most parsimonious model of the paternity assignments included genetic relatedness and colony size, we did not find strong evidence supporting their effect on fertilisation success. While intermediate genetic distances appeared to align with higher fertilisation success in some cases, this trend was not statistically significant. Likewise, and contrary to expectations, individual colony size also showed no statistically significant effect. Previous modelling exercises have highlighted mean population colony size as a major determinant of fertilisation success in broadcast-spawning corals, given established size-fecundity relationships (Hall & Hughes 1996, Álvarez-Noriega et al. 2016, Teo & Todd 2018, Ricardo et al. 2024b). However, while population-level mean colony size may influence fertilisation, individual colony size showed minimal impact within our experimental setup. Instead, we observed that distance and relative positioning concerning current direction were more influential factors.

Limited gamete mixing among nearby individuals at lower population densities restricts the potential for successful reproductive partnerships and thereby may act to constrain genetic diversity. Using empirically derived pairwise distances of parents from the experiment, simulations explored gametic mixing patterns across various colony densities under spatial distributions typical of A. hyacinthus reef environments incorporating both observed natural patterns and random distributions. These simulations indicated a relationship between colony densities and simulated gametic mixing, measured by successful pairwise crosses and the number of partners per individual (breeding units), with a marked increase observed at higher densities. This increased acceleration in mixing was observed above 0.01 colonies m^-2^, suggesting conditions that may facilitate broader gamete exchange, potentially resembling a well-mixed gamete suspension for this species. Conversely, at lower densities, these indicators of gametic mixing remained notably limited and relatively stable, while the number of isolated colonies increased. This suggests a lower density range below which effective gametic and, consequently, genetic mixing is limited.

While theoretical models assessing the retention of genetic diversity in ideal scenarios suggest that founding populations can retain a significant portion of source genetic diversity (Shearer et al. 2009), these often assume random mating (panmixia), minimum reproductive variance, and uniform gene flow across the population in subsequent generations (Ryman & Laikre 1991, Laikre et al. 2010). However, the reality of degraded reefs and small outplanted populations in the context of restoration, exacerbated by factors like fertilisation Allee effects, may violate these assumptions. Reduced colony density and subsequent spatial isolation of colonies limit effective gamete exchange among potential reproductive partners, resulting in substantially smaller effective breeding units. This fundamental constraint on effective gametic mixing reduces the effective population size (Ne), possibly increasing the risk of inbreeding depression over time, thereby eroding population genetic diversity and reducing fitness (Grupstra et al. 2024). Such conditions promote patterns like chaotic genetic patchiness and ‘sweepstakes reproductive success,’ where successful reproduction is highly localised and dominated by a few individuals or patches (Eldon et al. 2016, Cros et al. 2020, Barfield et al. 2023). Integrating evolutionary genetic modelling with these findings may help refine population density thresholds critical for restoration strategies aimed at sustaining diversity and promoting the spread of beneficial traits (DeFilippo et al. 2022, Humanes et al. 2024, Quigley 2024), though validation using pairwise parental distances from natural populations remains necessary.

These findings have implications for coral outplanting and restoration design, emphasising the importance of spatial arrangement and current direction to maximise reproductive success. Clustered outplants of corals will ultimately result in a greater likelihood of fertilisation success; indeed, high fertilisation rates were observed in the central patch of this study. However, even within the central cluster, some individuals did not participate in fertilisation, likely due to spawning asynchrony or because their offspring were not captured in the subsampled larvae. This, along with lower fertilisation observed in other studies (Ricardo et al. 2024b), highlights the need for outplant patches to be sufficiently large (>20 colonies) and dense to accommodate the possibility of some asynchronous spawning, less favourable hydrodynamic conditions, and to account for colony mortality after outplanting. Predicting current direction at the time of spawning remains challenging without site-specific hydrodynamic data, and its variability further complicates predictions. Maintaining a circular or radial outplanting design may help mitigate risk in the absence of such information. Furthermore, advancements in image-based survey methods may enhance the quantification of spatial population parameters, facilitating the identification of site-specific thresholds for successful fertilisation (Lechene et al. 2024, Remmers et al. 2024).

Our method of using fragments along the radial lines may hinder, though not entirely prevent, accurate assessment of fertilisation from the most distant colonies, as sperm from nearby colonies are more likely to reach and fertilise eggs sooner than those from distant sources, thereby potentially biasing the resulting offspring towards closer colonies. However, the low fertilisation success and attribution through paternity tests in almost all the samples from colonies at 30 m indicate that this was unlikely. Additionally, our design featured a greater number of close neighbour distances, and we acknowledge that our results may therefore have a bias to lower intercolonial distances than occurs in a natural population. Yet, recent work by Mumby et al. (2024) indicated that pairwise nearest neighbour (or intercolonial) distances best described observed fertilisation trends across a natural population in Palauden. Other studies have investigated relationships using proxies for adult densities (Coma & Lasker 1997, Levitan et al. 2004). However, density-fertilisation relationships are not always consistent across depths, species, and reproductive modes (Miller & Mundy 2005). Mumby et al. (2024) highlight the challenges with using ‘density’ as solely a population parameter to assess Allee effects, since clustering can cause density estimates to vary depending on the chosen spatial scale. These results are consistent with Levitan et al. (1992), who found that larger population sizes of the sea urchin Strongylocentrotus franciscanus led to higher fertilisation success, even when density was maintained constant. However, when area was controlled and density was allowed to vary, fertilisation success was influenced by changes in density. Spatially explicit simulation models have supported the use of population density as a relevant population parameter (Teo & Todd 2018, Ricardo et al. 2024b), likely because data are spatially averaged to the cell size inflating fine-scale mixing, and unlikely to result in very high or low fertilisation patchiness that nearest-neighbour parameters would determine. Ultimately, it is unlikely that a single population parameter or experimental design on Allee effects during fertilisation can address all these aspects, and multiple approaches combining survey, experimental and modelling exercises are needed (Levitan 1995).

Our study provides empirical evidence that fine-scale spatial arrangements of adult colonies on reefs are critical for maintaining reproductive success in broadcast spawning corals. As reefs continue to degrade, resulting in lower adult densities and altered spatial clustering, populations will increasingly approach or even exceed the minimum spatial thresholds for effective gametic mixing. These factors are likely to act alongside other processes that can reduce reproductive output (Briggs et al. 2024). Our results suggest that small, isolated patches of corals with low densities of colonies may face significant Allee effects, ultimately impeding natural recovery and resilience. These findings have implications for restoration strategies: efforts must prioritise not only increasing colony numbers but also ensuring optimal spatial configurations in spawning events to ensure fertilisation success and sustain genetic diversity. Integrating localised hydrodynamic data and tailored outplant designs may therefore be key to mitigating the cascading effects of reef degradation and securing the long-term viability of coral populations.

## Data availability statement

The datasets used for this study are deposited in the CSIRO Data Access Portal https://data.csiro.au/ and will be made fully available following publication.

## Acknowledgments

The work was conducted under CITES permit PW23-079 and PWS2023-AU-001443, Bureau of Marine Resources #100939-E, and Marine Research Permit RE-22-11. We thank R. Griffith-Mumby and A. Donovan for coordination support. We thank the Palau International Coral Research Center (PICRC) for research support. We thank Geory Mereb and other PICRC skippers for vessel support. We thank Anca Rusu for her assistance and advice with DNA extraction methods. This work was supported by the EcoRRAP subprogram (https://gbrrestoration.org/program/ecorrap/) that is part of the Reef Restoration and Adaptation Program (RRAP, https://gbrrestoration.org/). RRAP is funded by the partnership between the Australian Governments Reef Trust and the Great Barrier Reef Foundation. The funders had no role in study design, data collection and analysis, decision to publish, or preparation of the manuscript.

## Supplementary Material

Text S1. PCA revealed no clear separation of individuals into groups, with the first two principal components explaining only 7.6% and 6.9% of the total variance, respectively (Fig. S1). K-means clustering did not identify distinct groups, and the Elbow method indicated no strong inflection point, suggesting continuous genetic variation rather than discrete clusters. A PERMANOVA test revealed a significant difference between groups at K = 2 (p < 0.001), but the effect size was low (R^2^ = 0.067), suggesting that while some weak genetic differentiation exists, it explains only a small proportion of the overall variance. While ΔK was greatest at K = 2, STRUCTURE analysis exhibited low overall admixture, with most individuals predominantly assigned to a single cluster in the admixture model (data not shown). Mean LnP(K) values decreased slightly from K = 1 to K = 2, indicating that adding an additional cluster did not improve model fit, further supporting a lack of strong genetic structuring. Hardy-Weinberg equilibrium (HWE) tests revealed no loci remained significantly out of equilibrium after FDR correction, consistent with random mating and a panmictic population structure. Collectively, these results indicate weak genetic structuring, if any, with likely high connectivity across this small spatial scale leading to predominantly continuous genetic variation rather than discrete genetic groups.

**Fig. S1.**
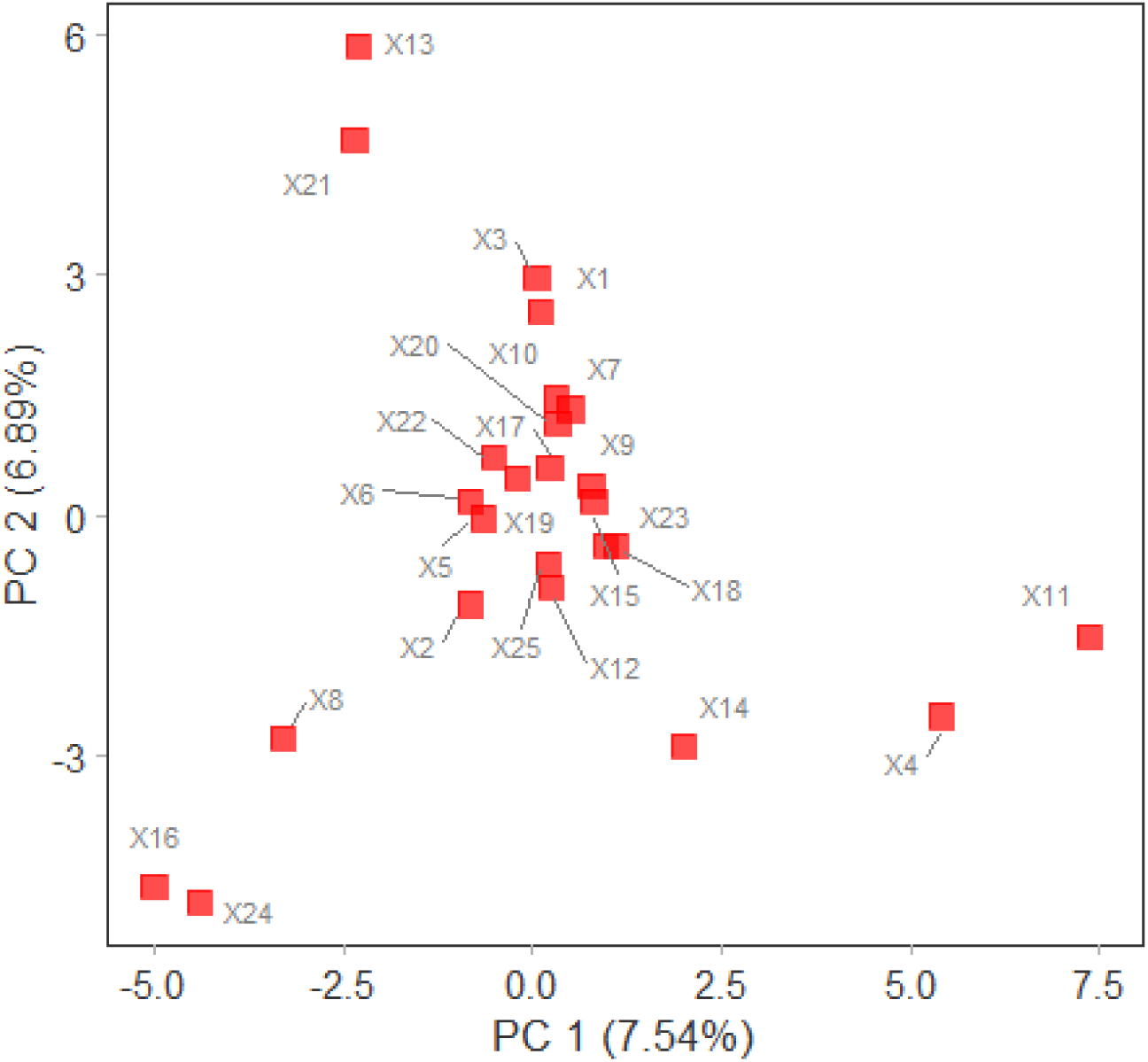
Principal Component Analysis (PCA) plot of distinct adult colonies of Acropora hyacinthus, showing ordination along the first two principal components.

**Fig. S2.**
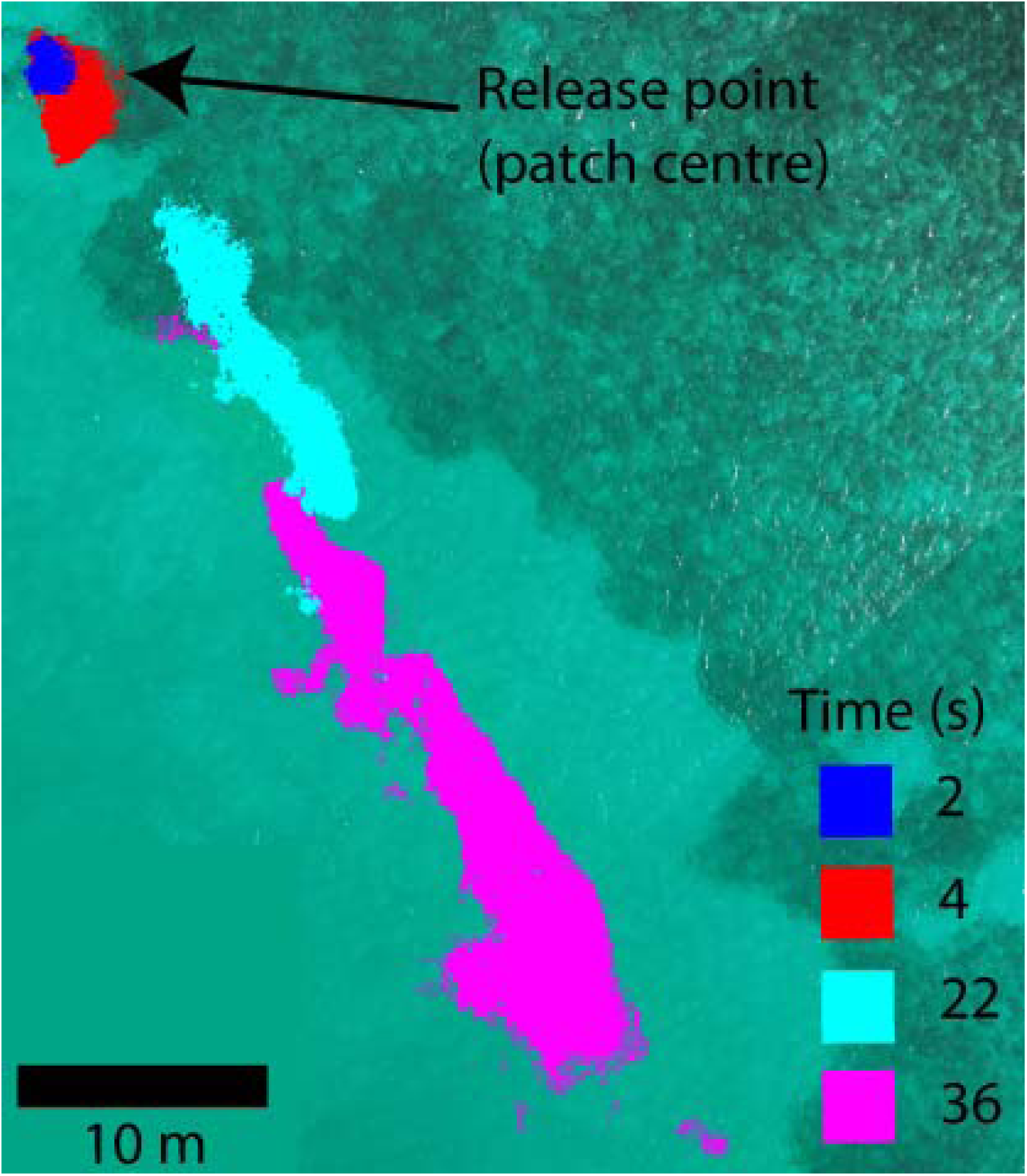
Movement of tracer dye over time (s) from the patch centre, recorded during a tidal cycle comparable to that of the field experiment.

**Fig. S3.**
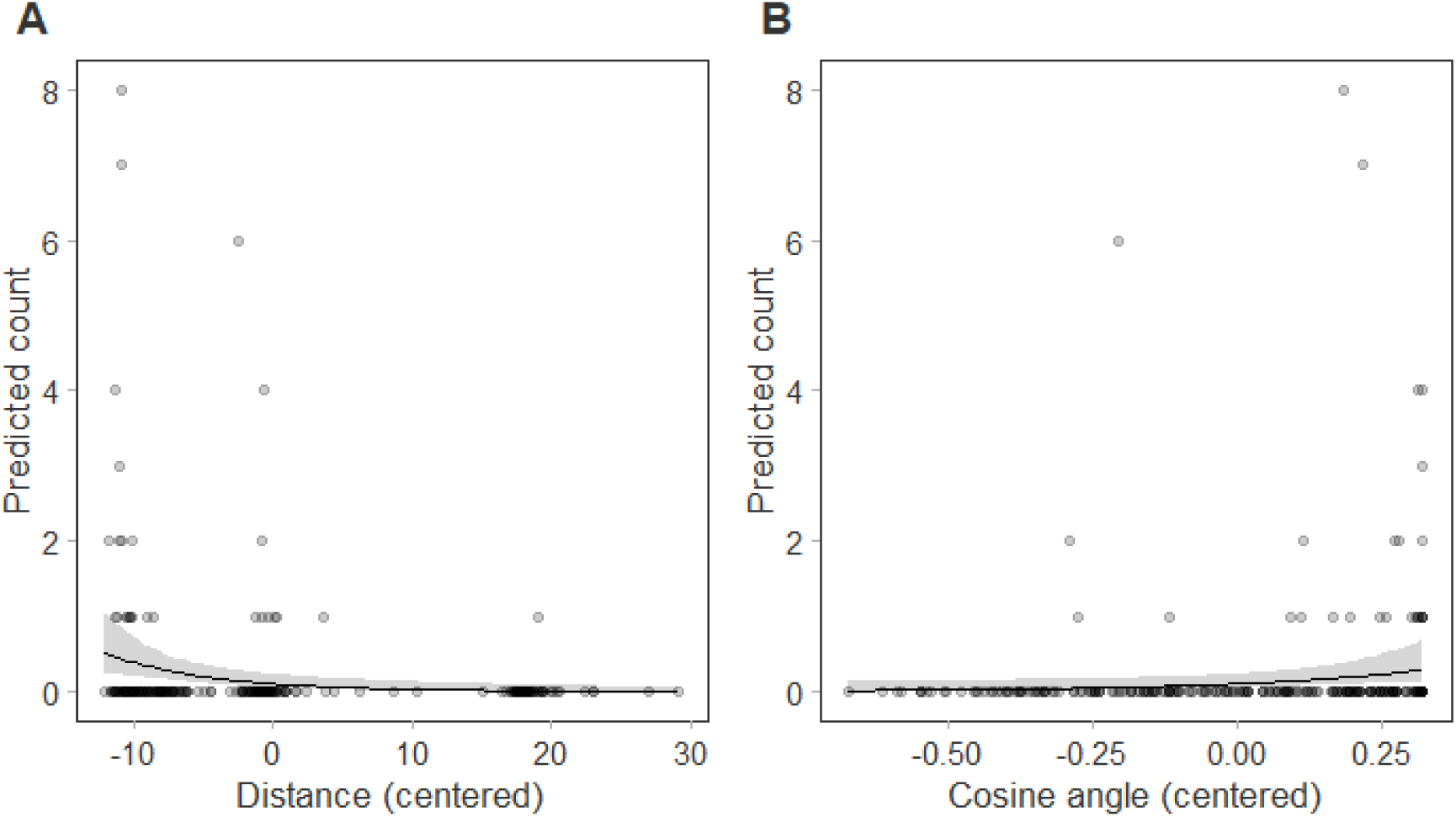
Effect plots of the predicted relationships between assigned parents and explanatory variables. Panels show the effects of (A) physical distance between colonies, (B) relative angular position of colonies to upstream/downstream angle, while holding other covariates at their mean values. Points represent raw data, solid lines show model predictions, and shaded areas represent 95% confidence intervals. Note that distance and angle values are mean-centred.

**Fig. S4.**
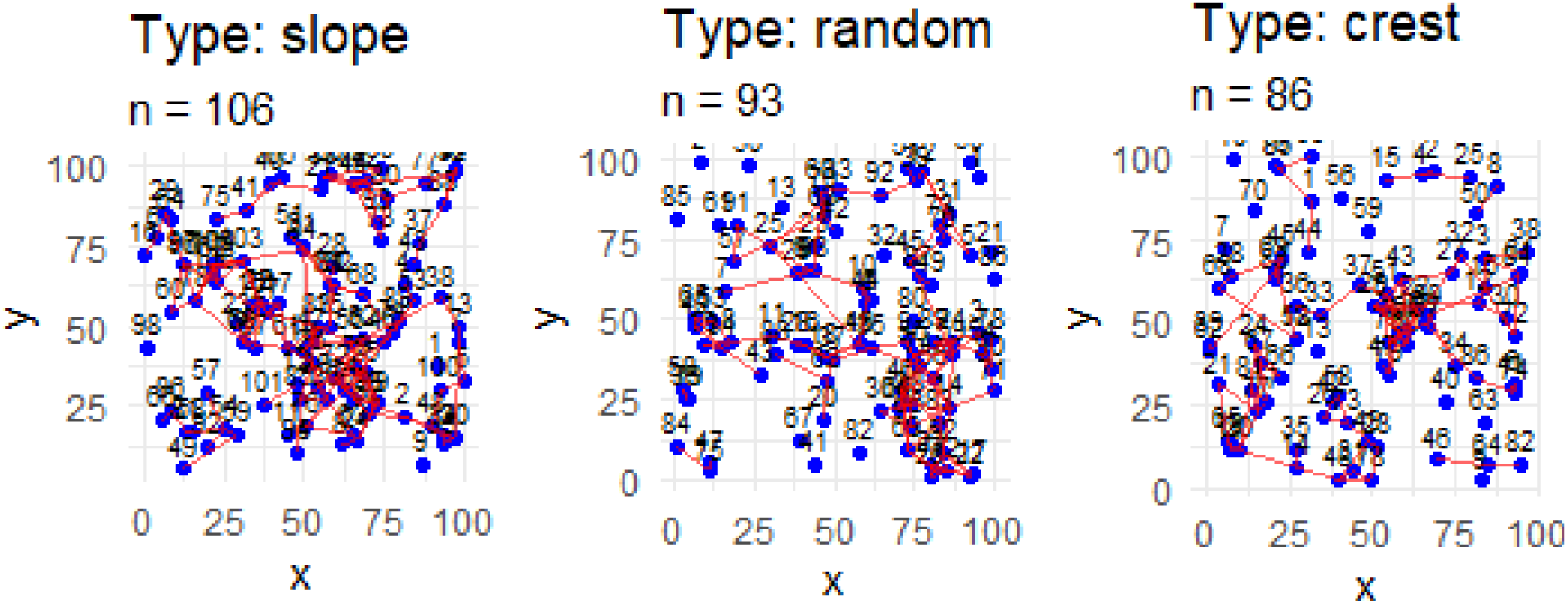
A representative example of simulated pairwise crosses at each reef spatial distribution (reef slope, random/Poisson, and reef crest) for the density = 0.01 colonies m^-2^ (100 colonies per hectare). Note that population size varies slightly between each run due to the random spatial arrangement.

## References

Álvarez□Noriega M, Baird AH, Dornelas M, Madin JS, Cumbo VR, Connolly SR (2016) Fecundity and the demographic strategies of coral morphologies. Ecology 97:3485–3493

Anderson EC, Garza JC (2006) The power of single-nucleotide polymorphisms for large-scale parentage inference. Genetics 172:2567–2582

Barfield S, Davies SW, Matz MV (2023) Evidence of sweepstakes reproductive success in a broadcast□spawning coral and its implications for coral metapopulation persistence. Mol Ecol 32:696–702

Barton K, Barton MK (2015) Package ‘mumin’. Version 1:439

Boström-Einarsson L, Babcock RC, Bayraktarov E, Ceccarelli D, Cook N, Ferse SC, Hancock B, Harrison P, Hein M, Shaver E (2020) Coral restoration–A systematic review of current methods, successes, failures and future directions. PloS one 15:e0226631

Briggs ND, Page CA, Giuliano C, Alessi C, Hoogenboom M, Bay LK, Randall CJ (2024) Dissecting coral recovery: bleaching reduces reproductive output in Acropora millepora. Coral Reefs 43:557–569

Buccheri E, Ricardo GF, Babcock RC, Mumby PJ, Doropoulos C (2023) Fertilisation kinetics among common Indo-Pacific broadcast spawning corals with distinct and shared functional traits. Coral Reefs 42:1351–1363

Coffroth MA, Lasker HR (1998) Larval paternity and male reproductive success of a broadcast-spawning gorgonian, Plexaura kuna. Mar Biol 131:329–337

Coma R, Lasker HR (1997) Effects of spatial distribution and reproductive biology on in situ fertilization rates of a broadcast-spawning invertebrate. The Biological Bulletin 193:20–29

Courchamp F, Clutton-Brock T, Grenfell B (1999) Inverse density dependence and the Allee effect. Trends Ecol Evol 14:405–410

Crimaldi JP (2012) The role of structured stirring and mixing on gamete dispersal and aggregation in broadcast spawning. The Journal of Experimental Biology 215:1031–1039

Cros A, Toonen R, Karl SA (2020) Is post-bleaching recovery of Acropora hyacinthus on Palau via spread of local kin groups? Coral Reefs 39:687–699

DeFilippo LB, McManus LC, Schindler DE, Pinsky ML, Colton MA, Fox HE, Tekwa E, Palumbi SR, Essington TE, Webster MM (2022) Assessing the potential for demographic restoration and assisted evolution to build climate resilience in coral reefs. Ecol Appl 32:e2650

dela Cruz DW, Harrison PL (2020) Optimising conditions for in vitro fertilization success of *Acropora tenuis*, A. millepora and Favites colemani corals in northwestern Philippines. J Exp Mar Biol Ecol 524:151286

Doropoulos C, Vons F, Elzinga J, Ter Hofstede R, Salee K, Van Koningsveld M, Babcock RC (2019) Testing industrial-scale coral restoration techniques: harvesting and culturing wild coral-spawn slicks. Frontiers in Marine Science 6:658

Dray S, Dufour A-B (2007) The ade4 package: implementing the duality diagram for ecologists. Journal of statistical software 22:1–20

Dubé CE, Boissin E, Mercière A, Planes S (2020) Parentage analyses identify local dispersal events and sibling aggregations in a natural population of Millepora hydrocorals, a free□spawning marine invertebrate. Mol Ecol 29:1508–1522

Eldon B, Riquet F, Yearsley J, Jollivet D, Broquet T (2016) Current hypotheses to explain genetic chaos under the sea. Current zoology 62:551–566

Evanno G, Regnaut S, Goudet J (2005) Detecting the number of clusters of individuals using the software STRUCTURE: a simulation study. Mol Ecol 14:2611–2620

Flanagan SP, Jones AG (2019) The future of parentage analysis: From microsatellites to SNPs and beyond. Mol Ecol 28:544–567

Frieler K, Meinshausen M, Golly A, Mengel M, Lebek K, Donner S, Hoegh-Guldberg O (2013) Limiting global warming to 2 C is unlikely to save most coral reefs. Nature Climate Change 3:165–170

Furukawa M, Kitanobo S, Ohki S, Teramoto MM, Hanahara N, Morita M (2024) Integrative taxonomic analyses reveal that rapid genetic divergence drives Acropora speciation. Mol Phylogen Evol:108063

Golbuu Y, Gouezo M, Kurihara H, Rehm L, Wolanski E (2016) Long-term isolation and local adaptation in Palau’s Nikko Bay help corals thrive in acidic waters. Coral Reefs 35:909–918

Gouezo M, Langlais C, Beardsley J, Roff G, Harrison PL, Thomson DP, Doropoulos C (2025) Going with the flow: Leveraging reef□scale hydrodynamics for upscaling larval□based restoration. Ecol Appl 35:e70020

Grupstra CG, Gómez-Corrales M, Fifer JE, Aichelman HE, Meyer-Kaiser KS, Prada C, Davies SW (2024) Integrating cryptic diversity into coral evolution, symbiosis and conservation. Nature Ecology & Evolution 8:622–636

Hall V, Hughes T (1996) Reproductive strategies of modular organisms: comparative studies of reef□building corals. Ecology 77:950–963

Harrison PL, Babcock RC, Bull GD, Oliver JK, Wallace CC, Willis BL (1984) Mass spawning in tropical reef corals. Science 223:1186–1189

Harrison XA (2015) A comparison of observation-level random effect and Beta-Binomial models for modelling overdispersion in Binomial data in ecology & evolution. PeerJ 3:e1114

Hartig F, Hartig MF (2017) Package ‘dharma’. R package 531:532

Humanes A, Lachs L, Beauchamp E, Bukurou L, Buzzoni D, Bythell J, Craggs JR, de la Torre Cerro R, Edwards AJ, Golbuu Y (2024) Selective breeding enhances coral heat tolerance to marine heatwaves. Nature Communications 15:8703

IPCC (2018) Impacts of 1.5° C of global warming on natural and human systems. In: Bindi M, Brown S, Camilloni I, Diedhiou A, Djalante R, Ebi K, Engelbrecht F, Guiot J, Hijioka Y, Mehrotra S (eds) IPCC: Geneva, Switzerland

Jenkins TL, Martinelli M, Ellis CD, Stevens JR (2024) Exploring reported population differences in Norway lobster (Nephrops norvegicus) in the Pomo Pits region of the Adriatic Sea using genome-wide markers. PeerJ 12:e17852

Jones OR, Wang J (2010) COLONY: a program for parentage and sibship inference from multilocus genotype data. Molecular ecology resources 10:551–555

Kalinowski ST, Taper ML, Marshall TC (2007) Revising how the computer program CERVUS accommodates genotyping error increases success in paternity assignment. Mol Ecol 16:1099–1106

Kitanobo S, Toshino S, Morita M (2022) Genetic variation in released gametes produces genetic diversity in the offspring of the broadcast spawning coral Acropora tenuis. Scientific Reports 12:5026

Knowlton N (2001) The future of coral reefs. Proceedings of the National Academy of Sciences 98:5419–5425

Kramer AM, Dennis B, Liebhold AM, Drake JM (2009) The evidence for Allee effects. Popul Ecol 51:341–354

Kurihara H, Watanabe A, Tsugi A, Mimura I, Hongo C, Kawai T, Reimer JD, Kimoto K, Gouezo M, Golbuu Y (2021) Potential local adaptation of corals at acidified and warmed Nikko Bay, Palau. Scientific reports 11:1–10

Ladd MC, Burkepile DE, Shantz AA (2019) Near□term impacts of coral restoration on target species, coral reef community structure, and ecological processes. Restor Ecol 27:1166–1176

Laikre L, Schwartz MK, Waples RS, Ryman N (2010) Compromising genetic diversity in the wild: unmonitored large-scale release of plants and animals. Trends Ecol Evol 25:520–529

Lasker HR, Gutiérrez-Rodríguez C, Bala K, Hannes A, Bilewitch JP (2008) Male reproductive success during spawning events of the octocoral Pseudopterogorgia elisabethae. Mar Ecol Prog Ser 367:153–161

Lechene MA, Figueira WF, Murray NJ, Aston EA, Gordon SE, Ferrari R (2024) Evaluating error sources to improve precision in the co-registration of underwater 3D models. Ecological Informatics 81:102632

Levitan DR (1995) The ecology of fertilization in free-spawning invertebrates. In: Ecology of marine invertebrate larvae. CRC Press123-156

Levitan DR, Fukami H, Jara J, Kline D, McGovern TM, McGhee KE, Swanson CA, Knowlton N (2004) Mechanisms of reproductive isolation among sympatric broadcast spawning corals of the *Montastraea annularis* species complex. Evolution 58:308–323

Levitan DR, Petersen C (1995) Sperm limitation in the sea. Trends Ecol Evol 10:228–231

Levitan DR, Sewell MA, Chia F-S (1992) How distribution and abundance influence fertilization success in the sea urchin *Strongylocentotus franciscanus*. Ecology 73:248–254

Magnusson A, Skaug H, Nielsen A, Berg C, Kristensen K, Maechler M, van Bentham K, Bolker B, Brooks M (2017) glmmTMB: generalized linear mixed models using template model builder. R package version 01 3

Martin TG, Wintle BA, Rhodes JR, Kuhnert PM, Field SA, Low□Choy SJ, Tyre AJ, Possingham HP (2005) Zero tolerance ecology: improving ecological inference by modelling the source of zero observations. Ecol Lett 8:1235–1246

Miller K, Mundy C (2005) In situ fertilisation success in the scleractinian coral Goniastrea favulus. Coral Reefs 24:313–317

Mumby PJ, Anthony KR (2015) Resilience metrics to inform ecosystem management under global change with application to coral reefs. Methods in Ecology and Evolution 6:1088–1096

Mumby PJ, Sartori G, Buccheri E, Alessi C, Allan H, Doropoulos C, Rengiil G, Ricardo G (2024) Allee effects limit coral fertilization success. P Natl Acad Sci USA 121

Nozawa Y, Isomura N, Fukami H (2015) Influence of sperm dilution and gamete contact time on the fertilization rate of scleractinian corals. Coral Reefs:1–8

Oksanen J, Kindt R, Legendre P, O’Hara B, Stevens MHH, Oksanen MJ, Suggests M (2007) The vegan package. Community ecology package 10:631–637

Oliver J, Willis B (1987) Coral-spawn slicks in the Great Barrier Reef: preliminary observations. Mar Biol 94:521–529

Oliver JK, Babcock RC (1992) Aspects of the fertilization ecology of broadcast spawning corals: sperm dilution effects and in situ measurements of fertilization. Biol Bull 183:409–417

Omori M, Fukami H, Kobinata H, Hatta M (2001) Significant drop of fertilization of *Acropora* corals in 1999: An after-effect of heavy coral bleaching? Limnol Oceanogr 46:704–706

Ortiz JC, Pears RJ, Beeden R, Dryden J, Wolff NH, Gomez Cabrera MdC, Mumby PJ (2021) Important ecosystem function, low redundancy and high vulnerability: The trifecta argument for protecting the Great Barrier Reef’s tabular Acropora. Conservation Letters:e12817

Premachandra H, Nguyen NH, Knibb W (2019) Effectiveness of SNPs for parentage and sibship assessment in polygamous yellowtail kingfish Seriola lalandi. Aquaculture 499:24–31

Quigley K (2024) Breeding and selecting corals resilient to global warming. Annual Review of Animal Biosciences 12:209–332

Remmers T, Boutros N, Wyatt M, Gordon S, Toor M, Roelfsema C, Fabricius K, Grech A, Lechene M, Ferrari R (2024) RapidBenthos: Automated segmentation and multi□view classification of coral reef communities from photogrammetric reconstruction. Methods in Ecology and Evolution

Ricardo GF, Doropoulos C, Babcock RC, Adam AAS, Buccheri E, Robledo N, Uribe Palomino J, Mumby PJ (2024a) Spawning asynchrony and mixed reproductive strategies in a common mass spawning coral. bioRxiv:2024.2012.2006.627154

Ricardo GF, Doropoulos C, Babcock RC, Buccheri E, Khalil A, Mumby PJ (2024b) Critical thresholds of adult patch density and spacing during coral fertilisation. Research Square

Riginos C, Popovic I, Meziere Z, Garcia V, Byrne I, Howitt SM, Ishida H, Bairos-Novak K, Humanes A, Scharfenstein H (2024) Cryptic species and hybridisation in corals: challenges and opportunities for conservation and restoration. Peer Community Journal 4

Ryman N, Laikre L (1991) Effects of supportive breeding on the genetically effective population size. Conserv Biol 5:325–329

Shearer T, Porto I, Zubillaga A (2009) Restoration of coral populations in light of genetic diversity estimates. Coral Reefs 28:727–733

Sheets EA, Warner PA, Palumbi SR (2018) Accurate population genetic measurements require cryptic species identification in corals. Coral Reefs 37:549–563

Suggett DJ, Camp EF, Edmondson J, Boström□Einarsson L, Ramler V, Lohr K, Patterson JT (2019) Optimizing return□on□effort for coral nursery and outplanting practices to aid restoration of the Great Barrier Reef. Restor Ecol 27:683–693

Svensson CJ, Jenkins SR, Hawkins SJ, Åberg P (2005) Population resistance to climate change: modelling the effects of low recruitment in open populations. Oecologia 142:117–126

Teo A, Todd PA (2018) Simulating the effects of colony density and intercolonial distance on fertilisation success in broadcast spawning scleractinian corals. Coral Reefs 37:891–900

Vollmer SV, Palumbi SR (2002) Hybridization and the evolution of reef coral diversity. Science 296:2023–2025

Warner PA, Willis BL, Van Oppen MJ (2016) Sperm dispersal distances estimated by parentage analysis in a brooding scleractinian coral. Mol Ecol 25:1398–1415

Willis BL, Babcock RC, Harrison PL, Wallace CC (1997) Experimental hybridization and breeding incompatibilities within the mating systems of mass spawning reef corals. Coral Reefs 16:S53–S65

Wolanski E, De Le Court M, Lambrechts J, Kingfsord M (2024) Mechanisms enabling the self-recruitment of passive larvae in the Great Barrier Reef. Estuar Coast Shelf Sci:108976

Wolanski E, Hamner WM (1988) Topographically controlled fronts in the ocean and their biological influence. Science 241:177–181

Wood S, Wood MS (2015) Package ‘mgcv’. R package version 1:729

Yund PO (2000) How severe is sperm limitation in natural populations of marine free-spawners? Trends Ecol Evol 15:10–13

Zayasu Y, Suzuki G (2019) Comparisons of population density and genetic diversity in artificial and wild populations of an arborescent coral, Acropora yongei: implications for the efficacy of “artificial spawning hotspots”. Restor Ecol 27:440–446

